# Compilation of 29-year *postmortem* examinations identifies major shifts in equine parasite prevalence from 2000 onwards

**DOI:** 10.1101/756254

**Authors:** G. Sallé, J. Guillot, J. Tapprest, N. Foucher, C. Sevin, C. Laugier

## Abstract

Horses are infected by a wide range of parasite species that form complex communities. Parasite control imposes significant constraints on parasite communities whose monitoring remains however difficult to track through time. *Postmortem* examination is a reliable method to quantify parasite communities. Here, we compiled 1,673 necropsy reports accumulated over 29 years, in the reference necropsy centre from Normandy (France). The burden of non-strongylid species was quantified and the presence of strongylid species was noted. Details of horse deworming history and the cause of death were registered. Building on these data, we investigated the temporal trend in non-strongylids epidemiology and we determined the contribution of parasites to the death of horses throughout the study period. Data analyses revealed the seasonal variations of non-strongylid parasite abundance and reduced worm burden in race horses. Beyond these observations, we found a shift in the species responsible for fatal parasitic infection from the year 2000 onward, whereby fatal cyathostominosis and *Parascaris* spp. infection have replaced death cases caused by *S. vulgaris* and tapeworms. Concomitant break in the temporal trend of parasite species prevalence was also found within a 10-year window (1998-2007) that has seen the rise of *Parascaris* spp. and the decline of both *Gasterophilus* spp. and tapeworms. A few cases of parasite persistence following deworming were identified that all occurred after 2000. Altogether, these findings provide insights into major shifts in non-strongylid parasite prevalence and abundance over the last 29 years. They also underscore the critical importance of *Parascaris* spp. in young equids.

## 1. Introduction

Horses harbour complex macroparasite communities along their digestive tract, encompassing among other *Gasterophilus* larval stages (bots), nematodes (mainly strongylids and ascarids) and tapeworms (Anoplocephalidae). The control of this vast parasite community has largely relied on the regular use of anthelmintic drugs. Following a few decades of treatments, parasitologists have reported alteration of strongylid communities: *Strongylus* spp. prevalence drastically decreased over time whereas cyathostomin infection has become a major issue (Herd, 1990; Love and Duncan, 1991). This shift in species importance was derived from independent scattered pieces of evidence in the field (Herd, 1990; Love and Duncan, 1991). It was also associated with the development of drug resistant cyathostomin populations across the world (Fischer et al., 2015; Sallé et al., 2017; Tzelos et al., 2017; Nielsen et al., 2018). Independent evidence of ivermectin resistant *Parascaris* spp. populations have also been accumulating in recent years (Lyons et al., 2008; Laugier et al., 2012; Martin et al., 2018). However, the quantification of parasitic community evolution through time remains difficult and limited reports have been made so far.

Necropsy is a reliable method for the diagnosis of infection with equine intestinal parasites, especially those which are not detected (immature or larval stages) or may be underestimated (tapeworms) by routine coproscopy (Lyons et al., 1981; Lyons et al., 1984; Proudman and Edwards, 1992; Rehbein et al., 2013). Moreover, large parasites are easily recovered by careful examination of bowel contents and intestinal mucosal surfaces. Data derived by specific *post-mortem* examinations are hence more definitive than those obtained from other methods of investigation. This technic provides an invaluable tool for measuring parasite abundance and prevalence (Lyons et al., 1981). Using this technique, Tolliver et al. reported observations on the composition of equine parasite communities and their respective prevalence over a 28-year period in Kentucky (Tolliver et al., 1987). This work is, to our knowledge, the most extensive time series to date but it was based on a subset of horses selected to have patent strongyle infection (Tolliver et al., 1987). A decade later, further reports suggested a decrease in the prevalence of *Gasterophilus* spp. but a steady rate of infection by *Parascaris* spp. in young horses (Lyons et al., 2000). Most recent report from the same region suggested an increased infection rate by bots and tapeworms but was in line with the decline of *Strongylus* spp. (Lyons et al., 2018).

To our knowledge, there is no other example of comprehensive longitudinal analysis of equine parasite communities in any other region than Kentucky. In France, only scarce data are currently available on prevalence and intensity of equine intestinal parasites and they mainly rely on coproscopy results (Laugier et al., 2012; Traversa et al., 2012). The equine necropsy unit from the French agency for food, environmental and occupational health safety (ANSES, France) has been performing around 300 necropsies a year since 1987, following the same procedure.

Here, we present the parasitological data recorded in 1,673 young horses examined during a 29-year period to establish overall infection pattern in relation to host and environmental factors. Using known deworming history, we identified likely cases of drug resistance, and we relied on histo-pathological conclusions to determine parasite species contribution to the death of examined horses. We also estimated changes in species prevalence and abundance over time.

## 2. Material and methods

### 2.1. Animals

Material consisted of 1,740 young equids (2 to 24 months old) from 758 stud farms. Animals were derived from an equine population of 8,564 equids that were submitted for routine necropsy at the ANSES laboratory for animal health in Normandy from January 1987 to December 2015. This source of dead horses is hence relatively unbiased toward drug resistance. It also offers both a snapshot of non-strongylid communities and an assessment of cyathostominosis and verminous arteritis in the same young equid population from Normandy. We restricted our analysis to this subset of the total population as they more often suffer parasitic infection, and to avoid heterogeneity in the data.

Foals younger than 2 months were excluded from the study because none of them harbored the parasites searched. Necropsies were usually performed within a few hours after death but some were delayed for periods up to 24 hours. About 90% of the equids examined were from Normandy, which is the leading horse breeding region in France with roughly 10,500 foal births per year.

For every animal, relevant metadata including age, sex, breed, date of death, original stud farm were recorded. Indications about the last anthelmintic treatment administered were also collected in most cases.

A few observations were discarded before analysis, including an individual whose age was unknown and 32 records collected from geldings (as it was not possible to estimate any sex effect with so few observations). In addition, 33 cases from a bankrupted stud farm and one horse from a farm where no anthelmintic drug were given were also removed from the dataset to avoid spurious signal linked to a lack of parasite management. In the end, 1,673 cases from 735 studs were retained for analysis.

Foals were generally born in spring thereby resulting in a collinear relationship between the age at necropsy and the month when the necropsy took place. To account for this structure in the data, age was rounded to the closest month and clustered into two categories, being either less than or 1-year-old (foal) or older (yearling). Month of necropsy was binned into seasonal categories: winter (January to March), spring (April to June), summer (July to September) and autumn (October to December).

### 2.2. Necropsy technique and parasitological procedures

This study builds on data collected during routine necropsy. As a result, fine-grained parasitological examinations could not be systematically carried out due to time and material limitations. Parasite count was therefore performed for non-strongylid large parasites as these species are relatively easy to detect upon visual inspection. In addition, routine examination also included a research of encysted cyathostomin larvae in the mucosa and submucosa of the large intestine and of migrating *Strongylus vulgaris* larvae in the major arteries of the gastro-intestinal tract.

Throughout the study period, all necropsies were performed using the same complete protocol (Rooney, 1970; Collobert, 1995) and were implemented by the same team. Particularly, evisceration of the different parts of the digestive tract was carried out according to the procedure described by Rooney (1970) and specific examinations were performed for parasite recovery.

During evisceration, the stomach, small intestine, cecum and ascending colon were isolated with ligatures. Then, every organ was opened with scissors and their content was collected separately, spread on large trays and examined grossly for parasites. The mucosal surfaces were gently flushed with tap water and visually inspected for attached parasites. The other parts of the digestive tract such as the pharynx and the oesophagus were also examined. Species were searched in the vicinity of their preferential niche. Therefore, special attention was paid to bots in the oral cavity, pharynx, oesophagus and stomach; tapeworms were looked for in the small intestine, ileocaecal junction, cecum and right ventral colon, whereas ascarids and pinworms were looked for in the small intestine and in the ascending and small colon respectively.

Parasite specimens recovered were identified as to family (Anoplocephalidae), subfamily (Cyathostominae), genus (*Gasterophilus, Parascaris, Strongylus*) or species (*Oxyuris equi*) according to their anatomical location and published keys and illustrations (Lichtenfels, 1975; Jacobs, 1986; Price and Stromberg, 1987). *Gasterophilus* spp larval stage and species were determined during two consecutive years only (from March 1990 to February 1992) using a dissecting microscope (x30) and following appropriate guidelines (Wells and Knipling, 1938). Regarding tapeworms, specimens recovered from the small intestine, caecum and right ventral colon were preserved separately in 10% formalin in order to be microscopically examined later for the purpose of specific identification (Euzéby, 1966; Lichtenfels, 1975).

Tapeworms recovered from the small intestine were all examined and identified. For tapeworms present in other intestinal segments, the following protocol was implemented: when less than 100 specimens were counted, all the worms were identified. In cases of heavier infection (more than 100 tapeworms), a 10% aliquot was examined.

The cranial mesenteric artery and its major branches were opened and evaluated for lesions secondary to the migration of *Strongylus vulgaris* larvae. Adherent thrombi and granulation tissue were removed by scraping the intimal surface; then the parasites were recovered by dissecting carefully all these fragments and counted.

A special procedure was applied to detect cyathostomin larvae. At each site of the large intestines where the presence of parietal larvae was suspected by careful visual inspection, a 10 cm^2^ fragment of the digestive wall was removed, examined by mural transillumination technique (Reinemeyer and Herd, 1986) and then dissected under a dissection microscope at 10 to 30X magnification to confirm the presence of cyathostomin larvae. The number of larvae per cm2 was recorded. This technic is less sensitive than artificial digestion and can only detect large developing larvae (late third stage larvae, fourth stage larvae) and will miss early third stage larvae (Chapman et al., 1999). Nevertheless, it was chosen as an optimum between material capacities and detection sensitivity of developing cyathostomins. Fourth stage larvae were searched for by visual inspection of the caeco-colic content, recovered and inspected under a microscope to confirm their developmental stage from morphological criteria, i.e. cup-shaped buccal capsule and no visible cuticle (Brianti et al., 2009).

### 2.3. Determination of the cause of death

The cause of death was determined according to horse clinical history (duration of the disease and evolution, clinical signs and results of laboratory tests), observed lesions and the epidemiological context. The same person was in charge of categorizing the cause of deaths throughout the study period, thereby making observations comparable across the years.

Parasites were declared as the most likely cause of death when parasite recovery was associated with the following lesions:

- *Parascaris sp*.: intestinal obstruction, intussusception or rupture, toxemia and allergic shock following treatment in heavily infected foals (more than 30 worms);
- Tapeworms: presence of at least one tapeworm associated with ileal, ileo-caecal, caeco-caecal and caeco-colic intussusception, thickening of the ileal wall with obstruction, paralytic ileus at the ileocaecal valve;
- Larval cyathostominosis was suspected in case of extensive typhlocolitis including mucosal congestion, oedema, ulceration and necrosis, along with the presence of numerous encysted larvae (more than 10 larvae per cm^2^) and or hundreds of emerged L4 larvae in the bowel content.
- Infection with *S. vulgaris* larvae was considered as the cause of death when arterial infarction and necrosis of a bowel segment was diagnosed and was associated with verminous arteritis and thromboembolism.

### 2.4. Statistical analyses

Statistical analyses were carried out with the R software v3.5 (R Core Team, 2016). Parasitological data were analyzed following a binary outcome, *i.e*. infected or not, or as a continuous trait that quantifies the severity of the infection. The binary trait was modeled using logistic regression and a binomial link function, while raw worm counts were assumed to follow a negative binomial distribution, which is common for over-dispersed count data. This latter assumption was confirmed using the fitdistrplus v.1.0-14 package (Delignette-Muller and Dutang, 2015) by visual inspections of scatterplots of observed quantiles against the theoretical quantiles from three distributions, *i.e*. normal, Poisson and negative binomial (supplementary Figure 1).

For both type of trait, models were built as the sum of known fixed effects, *i.e*. horse sex, breed (French trotter, Thoroughbred, miscellaneous), age class (older than one year of age or not), and the season at which the horse died. We also added a binary variable encoding the time period, *i.e*. before or after the observed break in species prevalence through time. The break in parasite prevalence occurring around the year 2000 was inferred after regression of their estimated prevalence upon the year, using the segmented package (Muggeo, 2017). This strategy was chosen to account for the temporal trends in species prevalence and abundance; more complex mixed models including year as a random effect did not provide precise estimates and were faced with convergence issues when combined with a negative binomial link function. Fixed effects were subsequently kept or discarded by an AIC-based variable selection using the *stepAIC()* function from the MASS package (Venables and Ripley, 2002). This procedure aims at minimizing the residual variance while avoiding model overfitting. Horse sex was never retained during the variable selection procedure.

The cause of death was registered and classified as a binary outcome, i.e. of parasitic origin or not, and regressed upon horse breed and the season at which the horse was examined using logistic regression. The prevalence of fatal parasite infection was regressed upon the year of examination to establish whether it varied significantly between 1987 and 2015.

Mean estimates of the logistic regressions were exponentiated to obtain the relative risk associated with each variable level.

Due to the very low prevalence of *Parascaris* spp. in yearlings, modelling of worm burden and prevalence for these species was performed on the foal data only (n = 1,174 out of the 1,673 available observations).

Any test with *P*-value below 5% was deemed significant.

## 3. Results

### 3.1. Overall infection pattern by non-strongylid species

Average non-strongylid parasite burden and prevalence were in a lower range of values (Figure 1). Only 14 horses harboured *O. equi* and this species was not considered further. Bots were recovered from 409 out of the 1,673 equids examined *post-mortem* (24.4% prevalence, 95% c.i.: 22% - 26%). The number of *Gasterophilus* spp. per infected equid ranged between 1 and 889 (mean = 65.03 ± 90.46 and median = 35) and these were found in the stomach in most cases (380 out the 385 cases with observations; 10 horses presented instar attached to the oesophagus and five horses had larvae attached to their pharynx). Species determination was performed between March 1990 and February 1992 on 4,650 larvae sampled from a subset of 153 horses. At that time, *G. intestinalis* was the most prevalent species (37.9%) followed by *G. nasalis* (12.4%) and *G. haemorrhoidalis* (0.7%). Tapeworms were found in 289 equids (17.2% prevalence, 95% c.i.: 15% - 19%), and were almost exclusively located in the caecum (n = 224 cases). *A. magna* and *P. mamillana* were recovered in 2 horses each, including a co-infection with *A. perfoliata* in both cases. *Parascaris sp*. was recovered from the small intestine of 207 foals (17.6% prevalence, 95% c.i.: 15.5% - 19.9%) with an average abundance of 95 worms recovered (ranging from 1 to 1605 individuals).

**Figure 1.**
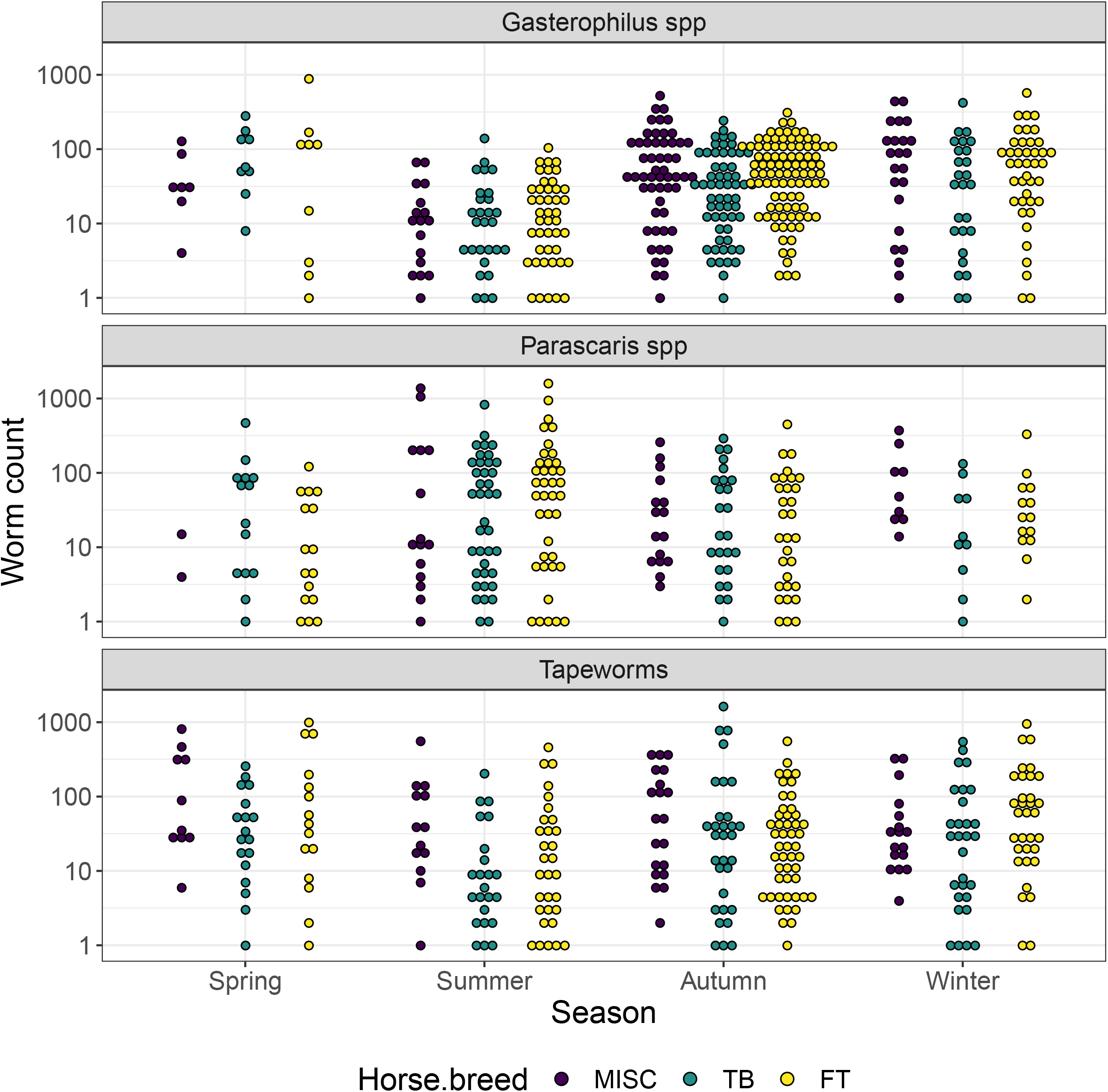
Non-strongylid species distribution across season and horse breed. The figure depicts the distribution of parasite burden measured in each breed type (MISC: Miscellaneous; TB: Thoroughbred; FT: French Trotter) and across seasons.

Co-infection by three non-strongylid species rarely occurred (n = 20), but 12.4% of the examined cases presented two non-strongylid species. In that latter case, parasites were twice as likely to be responsible for the death of the horse (14.4% of cases against 6.8% in the total population of cases).

The youngest foals with gastro-intestinal macroparasites were two months of age. Bots and *Parascaris* spp. were found in respectively 6 and 10 foals of that age with counts ranging from 1 to 34 and 1 to 75 individuals for bots and *Parascaris* nematodes respectively. The youngest foals infected with tapeworms were four months of age (n = 3) and harboured between 2 and 34 cestodes.

Parasite burden and prevalence followed seasonal fluctuations (Figure 1). *Parascaris* spp. were significantly more abundant in summer and autumn (averaged corrected burden of 34 ± 1.32 and 19.5 ± 1.42 nematodes/horse, *P* < 10^-4^ and 2 x 10^-3^), with a peak of infection in August. The same pattern was found for prevalence, whereby the highest frequency of infection was observed in autumn (odd ratio = 3.43, 95% c.i. = 2.12 - 5.57; *P* < 10^-4^). Bots and tapeworms hit their highest abundance later in the second half of the year, *i.e*. in autumn (*P* < 10^-4^) and winter (*P* = 0.018). During their respective most favourable season, bot and tapeworm prevalences was 14.67- (95% c.i.: 9.34 - 23.03) and 4.23-fold (95% c.i.: 2.77 - 6.46) as high as that observed in spring, respectively. Of note, tapeworms were more frequently found in yearlings than in foals (odd ratio = 2.54, 95% c.i. = 1.94 - 3.32) whereas horse age category neither contributed to bot burden variance nor to their prevalence variance.

Horse breeds were variously infected by non-strongylid parasites. Equids fell into three breed types: Thoroughbred (TB), French Trotter (FT) and miscellaneous (MISC) that encompassed French Saddlebreds (66.9%), other sport horses (6.6%), ponies (13.3%), Arabians (5.9) and draft horses (4.8%). Substantial variation was found in bot abundance and prevalence across the considered breed categories: Thoroughbred horses were significantly twice less likely to be infected by bots as miscellaneous horses (difference in relative risk = 0.45, 95% c.i. = 0.32 - 0.64; *P* < 10^-4^). In that case, parasite abundance was lower (8.58 ± 1.17 bots on average, *P* = 9.8 x 10^-3^) in Thoroughbred horses than for the two other breeds (13.46 ± 1.16 and 18.54 ± 1.29 bots on average for French trotter and miscellaneous horses respectively). Thoroughbred horses also displayed lower tapeworm burden on average (average burden of 11 ± 1.24 cestodes vs. 20 ± 1.24 and 39 ± 1.39 in French trotters and miscellaneous horses). However, their infection rate was not significantly different from that observed in other breeds (*χ*^2^ = 2.63, *d.f*. = 2; *P* = 0.27). No difference in *Parascaris* spp. (*χ*^2^ = 0.92, *d.f*. = 2; *P* = 0.63) infection rate was found between the three breed types considered.

### 3.2. A shift in parasite species causing the death of young horses through time

Out the 1,673 horses, most of them died spontaneously (n = 1347) whereas the remainder were euthanized by a veterinarian (n = 326). Overall, the cause of death was ascertained in 93.4% of horses (n =1563), suspected for 92 cases or remained unknown in 18 cases. Parasite were identified as being responsible for the death of 111 horses and highly suspected for 3 additional horses (Figure 2). Out of these, cyathostominosis was the most frequent cause of death (n = 38), followed by caeco-colic invagination caused by *Anoplocephala sp*. infection (n = 25). Thrombo-embolic disease caused by *S. vulgaris* (n = 22) and fatal *Parascaris* spp. infection (n = 19) were the main remaining causes of parasitic death.

**Figure 2.**
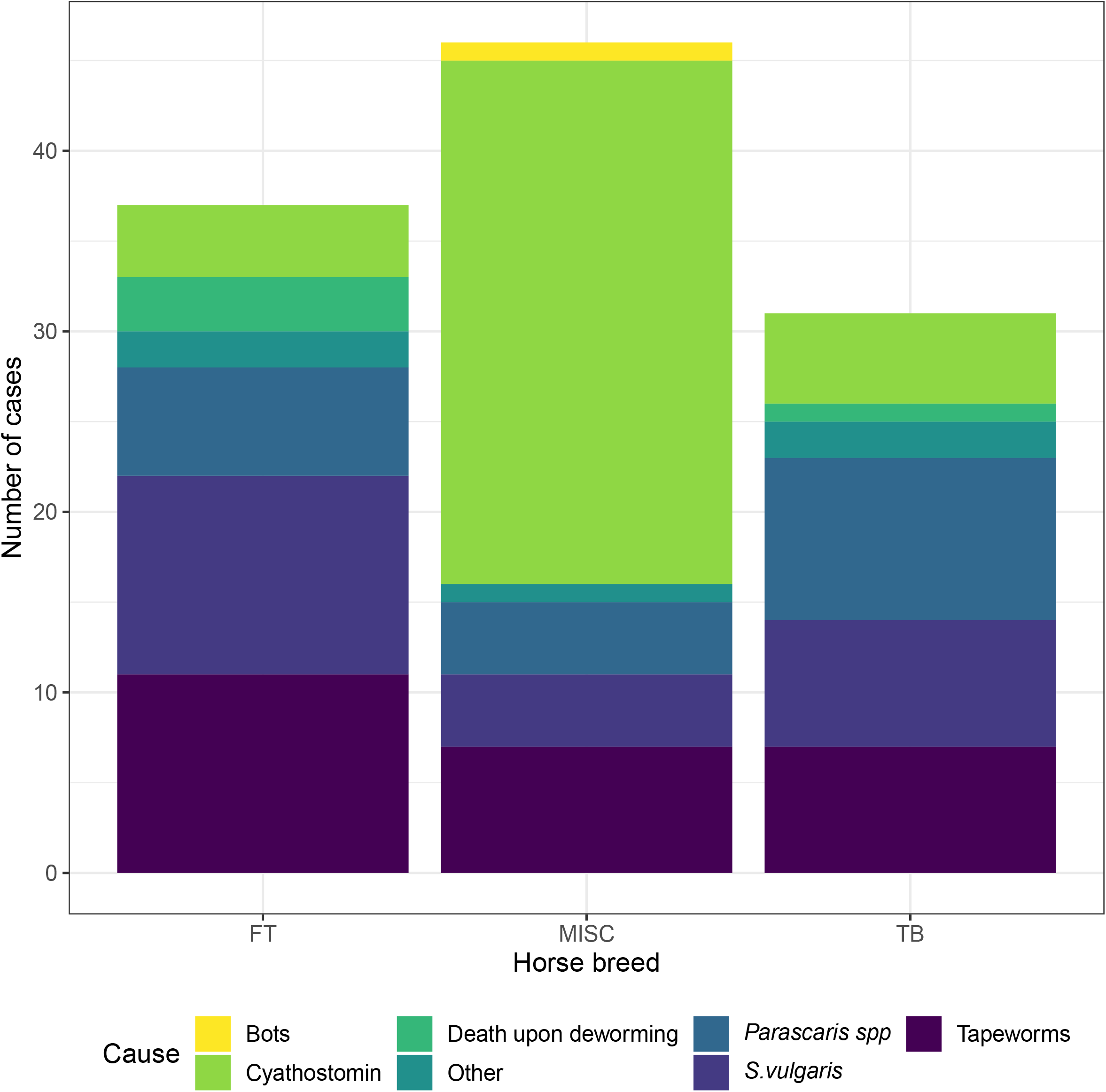
Parasites responsible for the death of young horses across breeds. The relative contribution of equine parasites to the death of young horses (114 cases) is plotted for each breed type considered (FT: French trotter; MISC: miscellaneous; TB: Thoroughbred). The figure highlights the higher contribution of cyathostominosis cases in the miscellaneous breed type.

The annual proportion of death caused by parasites remained relatively constant (6.5% ± 3.8% of total deaths) throughout the considered 29 years (*F*_1,27_ = 0.34; *P* = 0.56). It reached its highest in 2010 (18% of young horses necropsied) but was null in 2013. The relative risk of fatal parasitic infection was higher late in the year (4.1- and 6.4-fold increase in relative risk in autumn and winter respectively, *P* < 10^-4^). It was also significantly reduced in race horses (odd ratios of 0.37 and 0.39, *P* < 10^-4^ for both Thoroughbreds and French trotters respectively). In miscellaneous horses, cyathostominosis represented more than half of total deaths of parasitic origin (29 out of 38 cases) but this affection was less often seen in race horses (5 and 4 cases out of 31 Thoroughbreds and 37 French trotters respectively; Figure 2). French trotters were however more subject to fatal infection by tapeworms and *S. vulgaris* infection (Figure 2).

Of note, the yearly number of deaths caused by parasitic infection significantly increased after 2000 (2.53 ± 0.64 cases more, *P* = 10^-3^; supplementary Figure 2). A shift in the species responsible for the death of horses was also found from 2000 onward, whereby *S. vulgaris* and tapeworms have been progressively replaced by cyathostomins and *Parascaris* spp. in more recent times. *S. vulgaris* and tapeworms were responsible for 4.53 ± 0.92 (*P* < 10^-4^) more cases per year before 2000.

### 3.3. Persistence of gastro-intestinal helminths in recently dewormed horses

Complete deworming history including the date and class of the last anthelmintics used for deworming was available in 647 cases, 552 of which had been dewormed within the last 90 days. We found five cases (one French trotter, four Thoroughbred horses) of patent *Parascaris* spp. infection in foals that had been treated with ivermectin within the last 30 days before necropsy (4 to 22 days before death). These cases were noticed between 2004 and 2010. Two foals died because of *Parascaris* spp. mediated intestinal perforation, but the three others had non-parasitic causes of death.

*Parascaris* spp worms were found in foals treated with pyrantel two or six days before necropsy in 1999 and 2015 respectively. The former French trotter died from enterotoxaemia consecutive to deworming, while the latter Thoroughbred was euthanized because of a canon fracture.

A last case of patent *Parascaris sp*. infection was noticed on a 7.5 month-old Thoroughbred foal that had been drenched with fenbendazole four days before its death but harboured 54 worms.

An 18-month old Thoroughbred horse euthanized for a jaw lymphosarcoma in 2012, exhibited 836 *A. perfoliata* whereas he had been treated with a mixture of ivermectin and praziquantel 45 days before.

### 3.4. An increased prevalence of *Parascaris* spp. from 2008 onward

In relationship with the observed shift in species responsible for the death of young equids, we quantified the temporal variation of parasite prevalence and abundance across the 29-year period (Figure 3, supplementary Table 1). Breakpoints in non-strongylid prevalence were found to occur within a ten-year period around 2000, *i.e*. 1998, 2005 and 2007 for bots, tapeworms and *Parascaris* spp. respectively (Figure 3).

**Figure 3.**
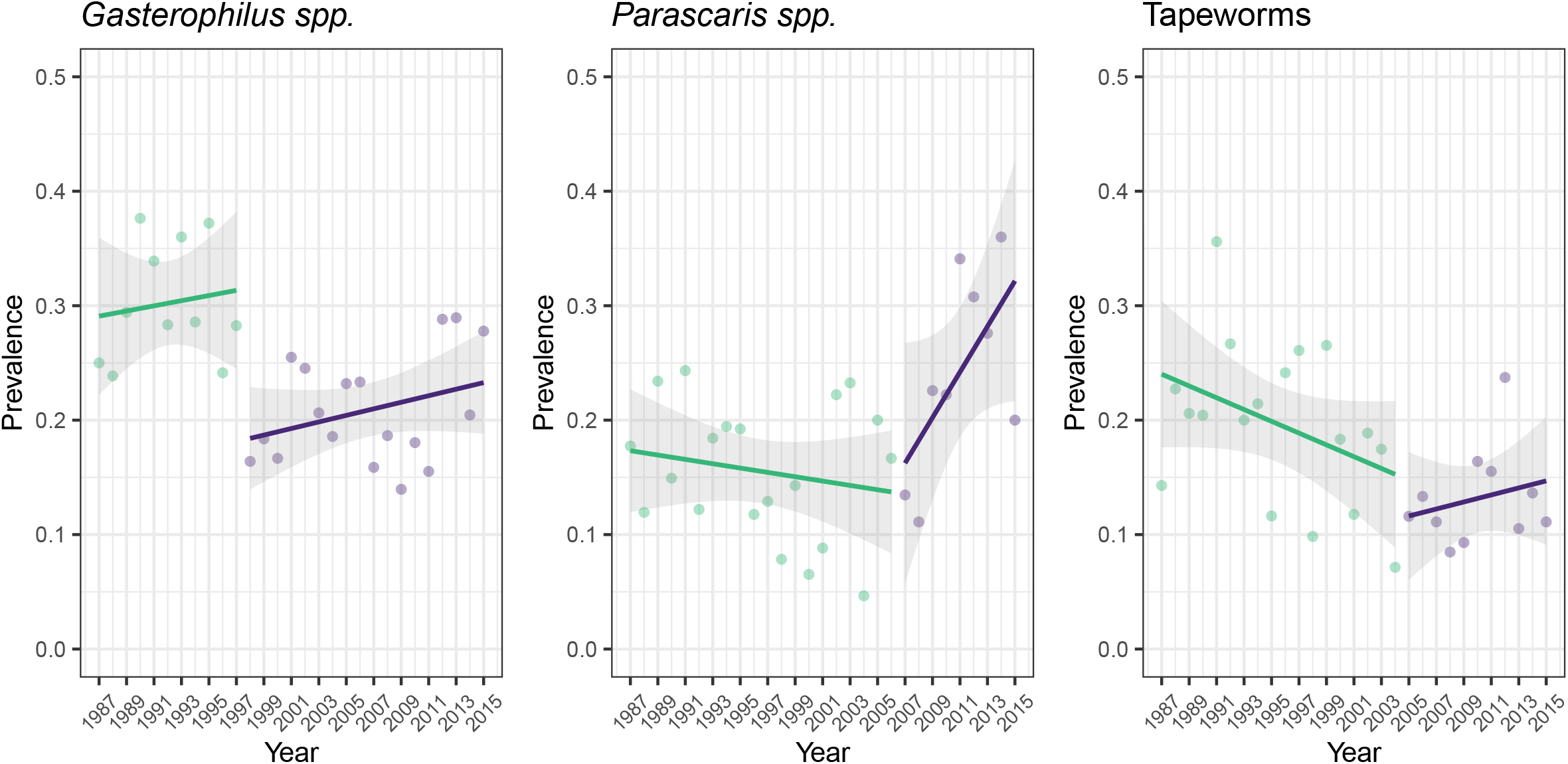
Temporal variation of non-strongylid parasite prevalence. The figure illustrates the breakpoints in parasite prevalence around the year 2000 for each of the three non-strongylid parasites considered, *i.e*. 1998, 2007 and 2005 for *Gasterophilus* spp., *Parascaris* spp. and tapeworms respectively. Points are coloured according to the considered time period, and the respective regression line is given with associated 95% confidence interval (shaded area).

This analysis revealed a 1.97-fold increase in the prevalence of *Parascaris* spp. infection after 2007 (95% c.i. = 1.41 - 2.75; *P* < 10^-4^). On average, 2.2 as many worms were observed in foals after 2007 relative to pre-2007 observations (*P* = 0.03). This suggests that following 2007, foals were significantly more often infected by *Parascaris* spp. and had increased worm loads.

An opposite pattern was found for bots and tapeworms (Figure 3). A break occurred in bots prevalence from 1998 onward, that resulted in a 1.35-fold (95% c.i.: 1.33 - 2.18; *P* < 10^-4^) reduction of its infection rate. This trend was also conserved for the abundance of bots found upon necropsy, with average count shifting from 17.8 ± 1.18 to 9.58 ± 1.15 after 1998 (*P* = 2.3 x 10^-3^). A similar significant reduction in the frequency of infection was found for tapeworm after 2005 (odds ratio = 0.62; 95% c.i.: 0.46 - 0.82; Figure 3), but their abundance was not significantly altered through time (*χ*^2^ = 0.12, *d.f*. = 1; *P* = 0.73).

## 4. Discussion

Our survey provides one of the most comprehensive long-term surveys of equine gastro-intestinal parasite dynamics. It is similar to a previous extensive report of 513 *postmortem* examinations performed between the mid-1950’s and 1983 in the USA (Tolliver et al., 1987). These horses had been however chosen because of their patent strongylid infection and the authors had limited information regarding their deworming history (Tolliver et al., 1987). This latter piece of information is difficult to obtain in field conditions and is often missing in *postmortem* examination (Lyons et al., 2000; Lyons et al., 2018) or abattoir surveys (Rehbein et al., 2013). In some studies, specific parasite species are searched for in a subset of individuals (Lyons et al., 2000). Here, we analyzed the long-term dynamics of parasite population in young horses, using the same examination protocol and the relevant background for each horse. The working subset of young animals reflected the diversity of equine production in Normandy. Indeed, horses were coming from 25% of the 2,981 stud-farms present in Basse-Normandy in 2014 (Anonymous, 2015). In addition, the diverse aetiologies underpinning the death of young equids suggest that these horses were not biased towards stud farms facing major issues in parasite control. A sampling bias remains however possible, as it is likely that all dead horses in the region were not sent for necropsy. The data collected on this subset of young horses highlighted a seasonal pattern in non-strongylid parasite abundance and prevalence. In agreement with previous reports from temperate areas, bots (Lyons et al., 1985; Price and Stromberg, 1987; Mfitilodze and Hutchinson, 1989; Lyons et al., 1994; Bucknell et al., 1995; Höglund et al., 1997; Lyons et al., 2000; Rehbein et al., 2013) and tapeworms (Benton and Lyons, 1994; Bucknell et al., 1995; Nilsson et al., 1995; Meana et al., 2005; Rehbein et al., 2013; Tomczuk et al., 2015) were more abundant and prevalent in autumn and winter seasons. This suggests that the subset of young equids, that were examined throughout the year, was a good proxy to investigate the regional parasite community dynamics. However, a peak of *Parascaris* spp. abundance was found in August and highest prevalence occurred in autumn. This slight seasonal disconnection between the occurrence of tapeworms and *Parascaris* spp may contribute to explain the limited extent of co-infection between the three non-strongylid species types. A similar seasonality was found in Northern Queensland (Australia), whereby *Parascaris* spp. infection was more prevalent in wetter months (Mfitilodze and Hutchinson, 1989). This finding is in contrast with multiple reports that did not find any evidence of a seasonal pattern (Lyons et al., 1994; Bucknell et al., 1995; Rehbein et al., 2013; Fabiani et al., 2016) and could result from the collinearity between the season when necropsies were performed and foal age. To this regard, the median foal age in August, when *Parascaris* spp. were the most abundant, was 4 months of age, which corroborates recent report (Fabiani et al., 2016).

Our prevalence estimates for bots and tapeworms were in the lower range of previously reported values, that varied between 15% (Lyons et al., 2000) to 94% (Tolliver et al., 1987) for bots and 30% (Mfitilodze and Hutchinson, 1989) to 80% for tapeworms (Benton and Lyons, 1994). This certainly reflects the important contribution of race horses to our dataset (84%), as these are usually subjected to intensive deworming programs. For instance, a 2013-survey across eight French Trotter studs revealed that foals were given eight anthelmintics a year (*Sallé et al*., unpublished observations).

Of note, a significant break in non-strongylid prevalences has occurred within a ten-year window ranging from 1998 to 2008, whereby *Parascaris* spp. arose in contrast to bots and tapeworms that suffered strong reduction in their respective prevalences. The sharp decline of tapeworm prevalence followed closely the release of praziquantel between 2001 and 2005, commercialized either alone (marketing authorization number FR/V/8052367 3/2001) or combined with ivermectin (marketing authorization number FR/V/1889939 3/2004) or with moxidectin (marketing authorization number FR/V/3281212 3/2005). The increased awareness of the association between tapeworm infection and clinical intestinal disease in horses in the 1990’s (Pearson et al., 1993; Proudman and Edwards, 1993; Proudman et al., 1998) has also certainly contributed to the implementation of a tapeworm-killing treatment in late fall or winter. At that time, tapeworm control relied on the off-label use of niclosamide (100 mg/kg) or a double dose of pyrantel embonate. On the contrary, the decrease in bot prevalence did not match the first release of ivermectin in 1983 (marketing authorization number FR/V/6151318 9/1983). Their decline was however lower than the 85% drop-off in *G. intestinalis* prevalence found between 1980 and 2000 in Kentucky (Lyons et al., 2000). Because species determination was performed during two years only, it was not possible to ascertain whether shifts in bots communities occurred. The observed decline remains hence difficult to explain with available data.

We also identified a significant shift in the species responsible for the death of young horses. Fatal tapeworm and *S. vulgaris* infections strongly declined after 2000, before a rise in *Parascaris* spp. and cyathostomin mediated deaths occurred. In the lack of farm management data or any climatic trend, a definitive explanation remains elusive. The decrease in fatal tapeworm infection is likely connected to its reduced prevalence starting in early 2000’s. The *S. vulgaris* decline has been reported since the 1990’s from various strands of evidence (Herd, 1990), although this is, to our knowledge, the first longitudinal quantification of this phenomenon. This observation is in strong contrast with recent observations from Scandinavian countries where increased prevalence of *S. vulgaris* was associated with evidence-based drenching regimens (Nielsen et al., 2012; Tydén et al., 2019). France, Denmark and Sweden fall under the same European regulation that imposes that anthelmintic drugs are delivered upon prescription by a veterinarian (Anonymous, 2001). However, the drug can be delivered without any coprological analysis in Sweden and France as opposed to the Danish setting (Anonymous, 1998). The limited uptake of evidence-based drug treatment in combination with the significant proportion of breeders buying anthelmintic drugs on their own in France (Sallé et al., 2017) is likely to explain the reduction in *S. vulgaris* prevalence.

Of note, cyathostominosis has remained the most frequent aetiology in death cases of parasitic origin. The miscellaneous horse category was particularly at risk in comparison to race horses. This higher incidence can arise from a poor control of cyathostomin populations or from a poor diagnostic and management of the horse condition. Recent survey on a limited number of premises in this region indicated that stud farms heavily relied on their veterinarians for the design of parasite management scheme but that it was not the case in riding schools (Sallé et al., 2017). This may contribute to increase cyathostomin prevalence outside stud farms. Non-professional horse owners or their veterinarians or both may also have a reduced awareness of cyathostominosis management which contribute to increase the incidence of fatal cases.

In the case of *Parascaris* spp., our observations suggest that a few cases of suboptimal drug efficacy occurred over the same time period. This is in line with other observations gathered from the same region (Laugier et al., 2012), from other European countries (Boersema et al., 2002; Schougaard and Nielsen, 2007; von Samson-Himmelstjerna et al., 2007; Näreaho et al., 2011; Martin et al., 2018) or from more distant areas, like in Australia (Beasley et al., 2015) or in the USA (Craig et al., 2007). This epidemiological context would hence suggest that the rise of *Parascaris*-mediated death might be linked to a decrease in anthelmintic efficacy.

As a conclusion, this compilation of *postmortem* examination over a 29-year period in a unique spatial entity, quantified major shifts in equine parasite communities that occurred within a 10-year window from early 2000 onwards. Observed patterns suggested that the release of macrocyclic lactones and praziquantel were major drivers of these shifts. The prevalence of fatal parasite infection remained constant through time, but fatal cyathostominosis cases have been increasing since the year 2000. This likely mirrors both a confusion with other causes of chronic diarrhoea by veterinarians in the field and a lack of awareness about drug resistance in cyathostomin populations resulting in a poor control of these populations. Worryingly, the rise of *Parascaris* spp. infection cases was concomitant with suboptimal anthelmintic efficacy cases that have appeared within the last decade. While additional education efforts among veterinarians and horse owners should contribute to dampen cyathostominosis cases, other strategies should be leveraged for the control of *Parascaris* spp. in foals.

## Acknowledgements

We are grateful to the Normandy Regional Council for its financial support to our equine necropsy unit since 1986. GS is a grateful recipient of an IFCE-Fonds Éperon grant (CYATHOMIX project).

---

**Supplementary Figure 1. Quantile-Quantile plot of non-strongylid parasite burden data** For every considered non-strongylid parasite (by row), empirical quantiles are plotted against theoretical quantiles from the normal (left column), Poisson (middle column) or negative binomial (right column) distributions. Any deviation from the black reference line is in favour of a mismatch between theoretical expectations associated with the chosen distribution and the observed data. For every parasite, data had their best fit against the negative binomial distribution.

**Supplementary Figure 2. Distribution of cyathostominosis cases throughout the study period** The number of fatal cyathostominosis cases seen over the 29-year period is plotted for each considered year.

