## Supplementary figures and images for "Compilation of 29-year *postmortem* examinations identifies major shifts in equine parasite prevalence from 2000 onwards"

### Supplementary Figure 1

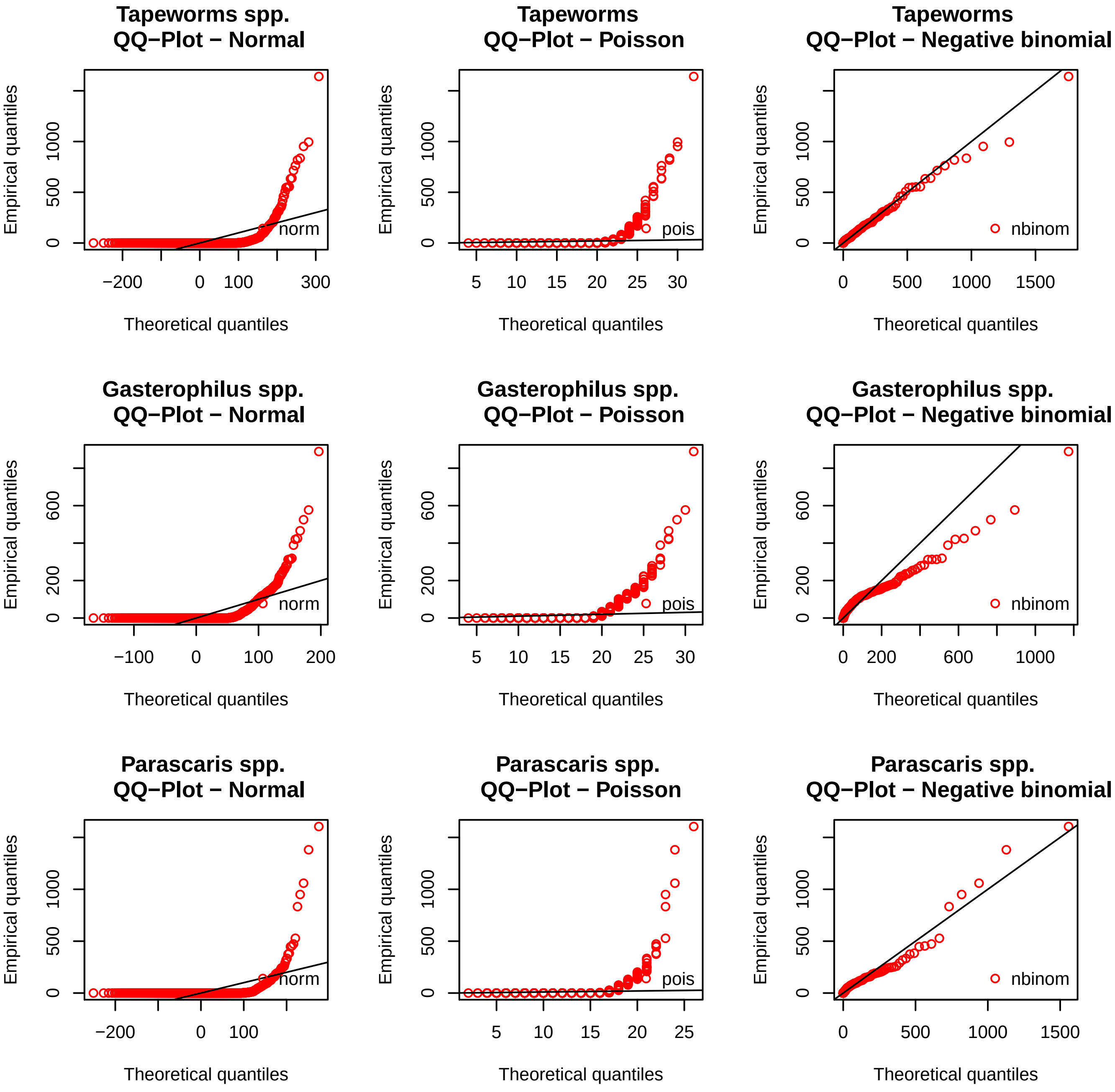

### Supplementary Figure 2

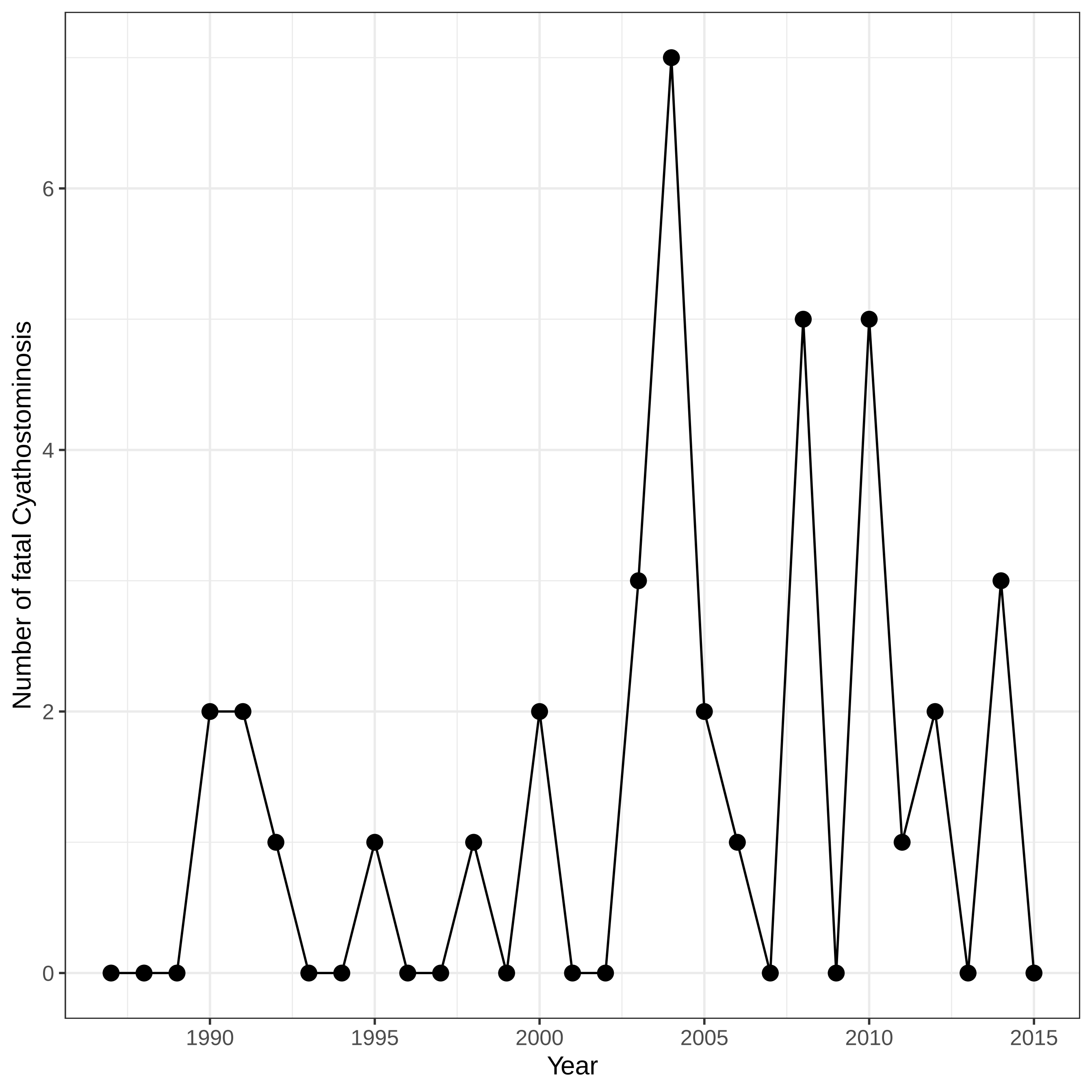
